# Selectivity matters: rules of thumb for management of plate-sized, sex-changing fish in the live reef food fish trade

**DOI:** 10.1101/098624

**Authors:** Holly K. Kindsvater, John D. Reynolds, Yvonne Sadovy de Mitcheson, Marc Mangel

## Abstract

Effective management of fisheries depends on the selectivity of different fishing methods, control of fishing effort, and the life history and mating system of the target species. For sex-changing species, it is unclear how the truncation of age structure or selection of specific size or age classes (by fishing for specific markets) affects population dynamics. We specifically address the consequences of plate-sized selectivity, whereby sub-mature, ‘plate-sized’ fish are preferred in the live reef food fish trade. We use an age-structured model to investigate the decline and recovery of populations fished with three different selectivity scenarios (asymptotic, dome-shaped, and plate-sized) applied to two sexual systems (female-first hermaphroditism and gonochorism). We parameterized our model with life-history data from Brown-marbled grouper (*Epinephelus fuscoguttatus*) and Napoleon fish (*Cheilinus undulatus*). ‘Plate-sized’ selectivity had the greatest negative effect on population trajectories, assuming accumulated fishing effort across ages was equal, while the relative effect of fishing on biomass was greatest with low natural mortality. Fishing such sex-changing species before maturation decreased egg production (and the spawning potential ratio) in two ways: average individual size decreased, and, assuming plasticity, females became males at a smaller size. Somatic growth rate affected biomass if selectivity was based on size-at-age because in slow growers, a smaller proportion of total biomass was vulnerable to fishing. We recommend fisheries avoid taking individuals near their maturation age, regardless of mating system, unless catch is tightly controlled. We also discuss the implications of fishing post-settlement individuals on population dynamics and offer practical management recommendations.

## Introduction

Fishing pressure can rapidly decrease population sizes, especially in large-bodied, longer-lived species with high commercial value (Colette et al. 2011, Sadovy de Mitcheson et al. 2013).In fisheries with limited abundance data, it is a major challenge to assess the status of a stock and make management recommendations (Dick and MacCall 2011, Carruthers et al. 2014). Furthermore, for many tropical species, even reliable catch data are lacking, requiring a form of triage in which life history traits are used to infer the species that could most benefit from management (Cheung et al. 2007, Kindsvater et al. 2016). There is still debate about which traits are most informative, and the best practices for management in data-poor situations (Gr<u>u</u>ss and Robinson 2014, Hordyk et al. 2015, Prince et al. 2015). An additional challenge arises when fisheries target small individuals of large species (Lee and Sadovy 1998, Reddy et al. 2013), as much of fisheries management has conventionally focused on the scenario where fishers prefer the largest fish available.

Groupers (Epinephelidae) are large, long-lived, high-value reef fishes (Coleman et al. 2000, Sadovy 2005, Rhodes et al. 2012, Craig et al. 2011, Sadovy de Mitcheson et al. 2013). They often form spawning aggregations, and many are female-first (protogynous) sex-changers. They are especially vulnerable to overfishing (Cheung and Sadovy 2005, Sadovy de Mitcheson and Liu 2008, Sadovy de Mitcheson 2016). Thus the life histories of groupers and other protogynous or aggregating fishes such as many wrasses and parrotfishes (Labridae) and the high market values of some commercially important species present a major challenge for managers and conservation biologists, especially in data-poor situations (Shepherd et al. 2013, ADMCF 2015). However, they also present some opportunities for management, such as temporal protection of spawning aggregations (Easter and White 2016, Sadovy de Mitcheson 2016). Prior work suggests that consideration of life-history traits could be especially informative for sex-changing and aggregating species (Molloy et al. 2009, Robinson and Samoilys 2013). Yet ow differences in selectivity among fisheries could interact with life-history traits is poorly understood.

In this paper, we develop a standard age- and size-structured model (Mangel 2006) parameterized for two sex-changing, aggregating and long-lived reef fish species. We compare the dynamics of a hypothetical separate-sex (gonochoristic) stock to those of female-first (protogynous) stocks. We explore how variation in growth (in particular in relation to size), natural mortality, and sexual system interact with fishery selectivity, which determines the size-or age-classes of fish that are caught (Maunder et al. 2014, Punt et al. 2014). We determine the steady-state spawning stock biomass and egg production (spawning potential) relative to the unfished population in each selectivity scenario. These metrics are commonly used in fisheries as biological reference points (Mangel et al. 2013, Kindsvater et al. 2016).

Our analysis of fishery selectivity differences is specifically motivated by the rapidly expanding and highly lucrative market for live reef food fish, which targets ‘plate-sized’ fish, typically between 20 and 40 cm (Reddy et al. 2013). The most valuable species in this trade include Brown-marbled grouper *E*. *fuscoguttatus*, Camouflage grouper *E*. *polyphekadion*, Leopard coral trout *Plectropomus leopardus*, Squaretail coral trout *P*. *areolatus*, and the Napoleon fish (or Humphead Wrasse) *Cheilinus undulatus* (Lee and Sadovy 1998, Sadovy 2005, Scales et al. 2007, Wu and Sadovy de Mitcheson 2016). However, there are very few data on the status of these fisheries. The species can be fished selectively when they are plate-sized. Alternatively, some are caught as juveniles and grown in captivity for months to years until they reach the preferred market size; this practice is known as capture-based aquaculture (CBA) (Lovatelli and Holthus 2008). For large-bodied species such as these, fishing targets sub-adult fish. Thus, we focus on the consequences of selectivity for stocks of the two largest species in the Asian-based live reef food fish trade. We compare the interaction between fishery selectivities and variation in life histories of these representative species. Although both species aggregate to spawn, this particular fishery focuses mostly on fish that have not reached reproductive maturity, rather than on adult, or spawning, fish.

Prior modeling of the population dynamics of female-first sex-changing fishes has been largely concerned with the consequences of possible sperm limitation for the productivity of the population (Alonzo and Mangel 2004, Heppell et al. 2006, Alonzo et al. 2008). Male depletion is undoubtedly cause for concern for species that have experienced dramatic changes in sex ratio, e.g. heavily female-biased populations of Gag grouper (*Mycteroperca microlepis*) (Heppell et al. 2006). However, biased sex ratios cannot explain why a number of sex-changing species have declined dramatically, since declines have happened even where sex ratios are near parity (e.g., in species with secondary males). Since the consequences of sperm limitation have been modelled (Alonzo and Mangel 2004, Heppell et al. 2006), we here focus on how fishing-induced changes in the size- and age-structure of the stock can affect biomass, egg production, and growth rate during recovery, as well as implications for management.

Using a Brown-marbled grouper-like life history as the first example, we address how size-selective fisheries that target sub-mature fish (specifically plate-sized fish) affect separate-sex stocks and contrast this with female-first stock dynamics. Although Brown-marbled grouper is also produced by full-cycle hatcheries, it is still taken extensively from the wild (Pears et al. 2006, Robinson and Samoilys 2013). We compare plate-sized selectivity (selectivity targeting 20-40 cm individuals), with asymptotic (targeting of largest fish available) and dome-shaped selectivity functions (targeting larger fish, but avoiding the largest sizes due to their behavior or gear selectivity). Dome-shaped selectivity often occurs in gillnet fisheries and is known to occur in data-rich fisheries such as North Sea cod (*Gadus morhua*, Maunder et al. 2014; Punt et al. 2014).

Our selectivity scenarios assume that *on average* fishing mortality across all ages is comparable, but distributed differently according to the fishery type or preference of the fishers. We chose the functions to contrast the markedly different selectivity pattern imposed in the live reef food fish trade with the selectivities commonly assumed for trawl or gillnet fisheries, or fisheries that are managed with a minimum length limit (Gwinn et al. 2015). We calculate biological reference points and rates of recovery after fishing has ceased for each selectivity type. We compare and contrast the Brown-marbled grouper with the Napoleon fish. In this wrasse species, males reach much greater maximum sizes than females, and small males are apparently rare. The consequences of this life history for management for the plate-sized fish favoured in this trade have never been explored, despite the fact that all Napoleon fish come from the wild and there is no commercial hatchery production.

We discuss the generality of our results and their implications for the practical management of live reef food fisheries targeting plate-sized fish. We also address fisheries that target small juveniles (to be grown to the target size in captivity), which differ from wild-capture fisheries targeting plate-sized fish. The sustainability of these fisheries depends on the change in natural mortality rate between size at settlement (typically < 5 cm) and larger fish, because fish populations experience a bottleneck around the age of settlement.

## Methods

We extend a commonly used age-structured population model to understand the potential for differences between the dynamics of female-first and separate-sex stocks (e.g., Haddon 2001, Alonzo and Mangel 2005, Mangel 2006). We conducted all programming in R (R Core Team, 2015). To follow the dynamics of biomass, we calculate size from age in the model. The size-at-age relationship is determined by our assumptions about individual growth (i.e., the von Bertalanffy growth function; see below). Growth underpins important life-history traits such as maturation, fecundity, and sex change, as well as natural and fishing mortality.

We first parameterized our model with life-history data for Brown-marbled grouper (Pears et al. 2006, Cushion 2010, Robinson and Samoilys 2013) (Table 1). We varied each parameter to determine the sensitivity of our results to variation in growth, maturation rules (e.g., age- vs. size-dependence), sex determination, natural mortality, recruitment, fishing mortality, and selectivity. We compared the model results with and without sex change. We then compared these results with predictions from models parameterized for the Napoleon fish (Table 1), for which we only modeled a sex-changing life history. This fish reaches a much larger body size but has similar sizes at maturation and sex change as Brown-marbled grouper. All females that will undergo sex change appear to transition to males by 80 cm, and small males are rarely documented (Sadovy et al 2007), so in this example we assumed the sex change function is a fixed function of size, although in sensitivity analyses we varied the degree of plasticity in sex change, as explained below.

### Age-structured model description

We modeled the dynamics of a population in age *a* and time *t* with overlapping generations. The population grows through recruitment to the first age class. We represent the number of females in this class as *N* (0, *t*). Assuming a constant sex ratio at birth and identical mortality, the number of males *N_m_*(*a, t*) ∝ *N*(*a, t*). The population declines as a function of natural mortality *M(a)* and fishing mortality *F*(*a,t*), which are both potentially functions of individual size. For *a > 0* the dynamics are

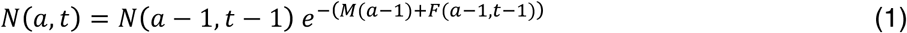

All individuals recruit to the population model at the same minimum size *L*(1). From then on, individuals grow according to the von Bertalanffy growth equation for length *L(a)* at age *a* (Fig. 1; Mangel 2006)

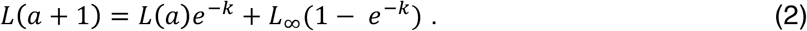

The parameter *k* determines how fast the fish grow to asymptotic size *L*_∞_, which we interpret as the maximum size possible. In our model of Brown-marbled grouper life histories, we assume that males and females have similar growth rates. However, for Napoleon fish, we assume that once sex change occurred, males grow faster than females. This is based on the observation of Choat et al. (2006) that protogynous males grow approximately twice as fast as females.

**Table 1.** Specific life history parameters for Brown-marbled grouper are from Pears et al. (2006). Parameters for Napoleon fish are from Choat et al. (2006), with the exception of the von Bertalanffy growth function parameters, which are from Sadovy et al. (2007). All other parameters are from Alonzo and Mangel (2005) or were varied in sensitivity analyses to determine if they affect results. The maturation and sex change ogives were adjusted to reflect the differences in observed patterns between the two species (e.g., small males are very rare in Napoleon fish, so we assumed a relatively shallow maturation ogive but a steep sex change ogive relative to Brown-marbled grouper).

| Parameter | Interpretation | Value for Brown-marbled grouper example | Value for Napoleon fish example |
| --- | --- | --- | --- |
| **SIMULATION PARAMETERS** |  |  |  |
| $t$ | Number of years in our population simulation. | 400 | 800 |
| $T_{\text{fishing}}$ | Time when fishing mortality begins | 150 | 500 |
| $T_{\text{recovery}}$ | Time when fishing mortality ends | 250 | 650 |
| **LIFE HISTORY PARAMETERS** |  |  |  |
| $M$ | Natural mortality rate: a function of age $a$, size, or constant | Varies from 0.1 to 0.2 | 0.1 |
| $A_{\text{max}}$ | Maximum age in years (after this age we no longer track individual fates) | 40 | 30 |
| $L_{\infty}$ | Asymptotic size (total length in cm) in the von Bertalanffy growth function | 80 | 91.5 for females;<br>168.4 for males |
| $k$ | Growth coefficient in the von Bertalanffy growth function | 0.16; sensitivity analyses varied from 0.1 to 0.2 | 0.13 for females;<br>0.06 for males |
| $L(a)$ | Length-at-age; $L(1)$ is the initial size at recruitment | 4 (when $a = 1$) | 4 (when $a = 1$) |
| $L_{\text{mat}}$ | Length at which 50% of females mature; | 45 | 45 |
| $q$ | Shape parameter that determines the slope of the maturation ogive | 0.7 | 0.1 |
| $\nu$ | Scale parameter relating length to body mass (value is arbitrary) | 1 | 1 |
| $\omega$ | Shape parameter relating length to body mass | 3 | 3 |
| $c$ | Scale parameter relating body mass to egg production (value is arbitrary) | 6 | 6 |
| $b$ | Exponent relating body mass to egg production (value is arbitrary) | 2 | 2 |
| $\rho$ | Shape parameter that determines the slope of the sex change ogive | 0.08 | 0.3 |
| $L_c$ | Size at which 50% of individuals have changed sex | 75 | 68 |
| $\Delta L$ | Discounting parameter; modulates the length at sex change relative to average lengths of the population | 27 | Varied from 0 to 20 |
| **RECRUITMENT PARAMETERS** |  |  |  |
| $\alpha$ | Parameter of the Beverton-Holt recruitment function; determines steepness near origin | 0.05 | 0.05 |
| $\beta$ | Parameter of the Beverton-Holt recruitment function | $1 \times 10^{-5}$ | $1 \times 10^{-5}$ |
| $\tau$ | Time ($\tau < 1$) spent in the larval stage prior to recruitment (Eq. 13) | - | - |
| **EXPLOITATION PARAMETERS** |  |  |  |
| $F_{\text{max}}$ | Maximum fishing mortality rate | Varies from 0.1 to 0.5 | Varies from 0.1 to 0.5 |
| $s(a)$ | Selectivity of the fishing gear for each age, can also depend on size $L(a)$ | Table 2 | Table 2 |
| **POPULATION METRIC** |  |  |  |
| $\lambda_{\text{rec}}$ | Growth rate of a recovering population after fishing ceases (after $T_{\text{recovery}}$; see Fig. 3) | - | - |

We assume mass at age *W(a)* and length *L(a)* are described by an allometric function

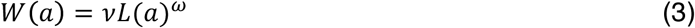

where *v* and *ω* are scale and shape parameters, respectively, which can be estimated from data (Alonzo and Mangel 2005).

**Figure 1.**
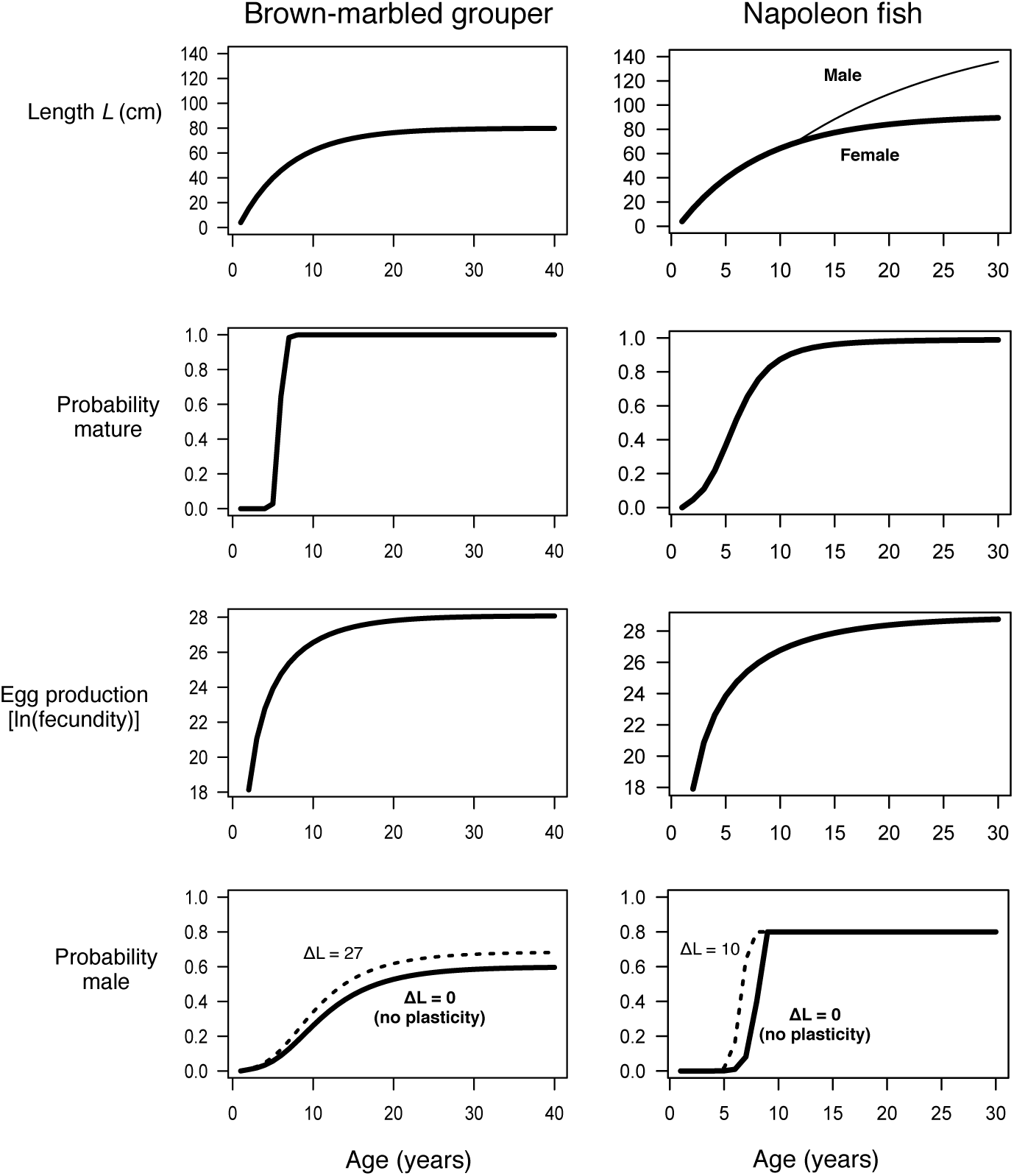
(a,b) Age-dependent length, (c,d) maturation, (e,f) fecundity and (g,h) sex-change functions for contrasting life histories of Brown-marbled grouper (left column) and Napoleon fish (right column). The latter species is larger overall, but matures earlier. In both cases, as maturation depends on size, and sex change depends on individual size and average size of the population, these functions will vary with somatic growth rate. Dashed curves in (g,h) represent changes in the sex change functions that arise as fishing reduces the mean length of individuals in the population (here shown for *M* = 0.1, asymptotic selectivity and *F_max_* = 0.3 for both species). Parameters are in Table 1.

### Reproduction

We assume the probability that an individual is mature *p_mat_* is a function of length-at-age (Figure 1)

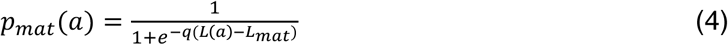

where *q* determines the steepness of the function and *L_mat_* determines the age at which half of the fish are mature. When *q* is large, maturation is knife-edged, in the sense that virtually all individuals below *L_mat_* are immature and those above are mature. When *q* is small, there is some probability of both immature and mature individuals at each size/age.

For simplicity, we assume that egg quality does not vary with female size. Egg production *E(t)* is determined by the number of females of age *a* at a given time *N(a,t)*, the probability females are mature *p_mat_(a)*, their biomass *W(a)*, their mass-specific fecundity, denoted by *c*, and an allometric parameter *b* that relates female mass and fecundity

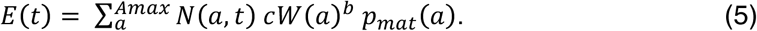

### Sex change

In monandric protogynous species (with no secondary males), individuals are born as females and turn into males as they grow larger. The probability that a female of age *a* and length *L(a)* becomes a male, *p_male_(a)*, is a function of body length-at-age

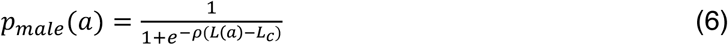

where *L_c_*. is the length at sex change at which half of the individuals are male. The length at sex change can be fixed, or it can be a function of the mean length of individuals in the population *L̅(t)* at that time, such that *L_c_(t)* = *L̅(t)* + Δ*L* (where Δ*L* offsets represents any difference between the population mean length and the local group size mean length (Alonzo and Mangel 2005); thus modulating the degree of the plastic response to changes in size-structure. In this case the probability of sex change will be a function of both *a* and *t*, as well as the social environment. Eq. 6 allows for the possibility that some females do not change sex despite approaching the maximum size (*cf* Pears et al. 2006, Choat et al. 2006).

Sex-changing fishes can vary in the degree of plasticity of the sex change function, so we model two example sex change functions. In our first example, females change sex according to their length relative to the population or group average: as the average length in the population decreases, the probability of becoming male increases for smaller individuals. In the second example (based on Napoleon fish) we assume that the sex change function responds less, or not at all, as the average length of the population decreases (i.e. there is less plasticity associated with age/size of sex change, although nothing is known of the factors that control sex change in this species; see the lower panels of Figure 1). We also assume that the probability of switching to male sizes once the female is greater than the sex change threshold is never greater than 0.8, because in Napoleon fish some older females evidently never change sex (Choat et al 2006).

In all female-first examples, we represent egg production as *Eh(t)*. In analogy to Eq. 5,

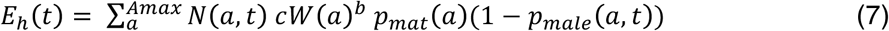

### Recruitment

The number of recruits, *N*(0, *t*), is determined by the number of eggs *E* produced by spawning female biomass in the previous time, as well as the density-dependent recruitment function, which we assume to follow the Beverton-Holt Stock Recruitment Relationship (SRR) (Beverton and Holt 1957, Mangel 2006, Mangel et al. 2013)

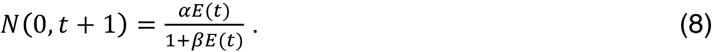

This determines the size of the year class that recruits to the population model (see Eq. 1). When *t* = l, we assumed there were 100 recruits seeding the population. In the discussion we adapt the recruitment function to show how capture of juveniles at settlement (when density-dependent mortality may substantially decrease juvenile numbers) could affect population dynamics (Mous et al. 2006).

### Mortality

For the majority of our analyses, we assume that individuals taken after settlement experience a fixed rate of natural mortality *M*, regardless of individual size. In sensitivity analyses, we also considered the case where natural mortality decreases for larger fishes (Lorenzen 2008, Brodziak et al 2009). However, changes in natural mortality rate with age did not strongly influence our results (see Results; Figs. S1-S3). We also assessed the effects of variation in somatic growth on biomass, which can be modeled by changing either *k* or *L*_∞_, making one or both smaller as biomass of the population increases (Munch et al 2005). We found that changing both parameters had similar effects, so we will discuss variation in the growth rate coefficient *k* while holding *L*_∞;_ constant.

Figure 2 shows the three selectivity functions *s(a)* that we considered: I) Asymptotic selectivity that is biased towards the largest individuals; II) dome-shaped selectivity, in which there is some preference for larger fish, but the very largest fish are not selected, e.g., due to gear selectivity or fishery type or behavior, such as habitat preference; and III) plate-sized selectivity, in which there is a target size which is approximately the size of a dinner plate. After growing past this length, the probability a fish is captured by the fishery is low (Fig. 2). For example, the desired size of Napoleon fish can be visually selected by snorkelers or divers who use a chemical to target specific individuals.

**Figure 2.**
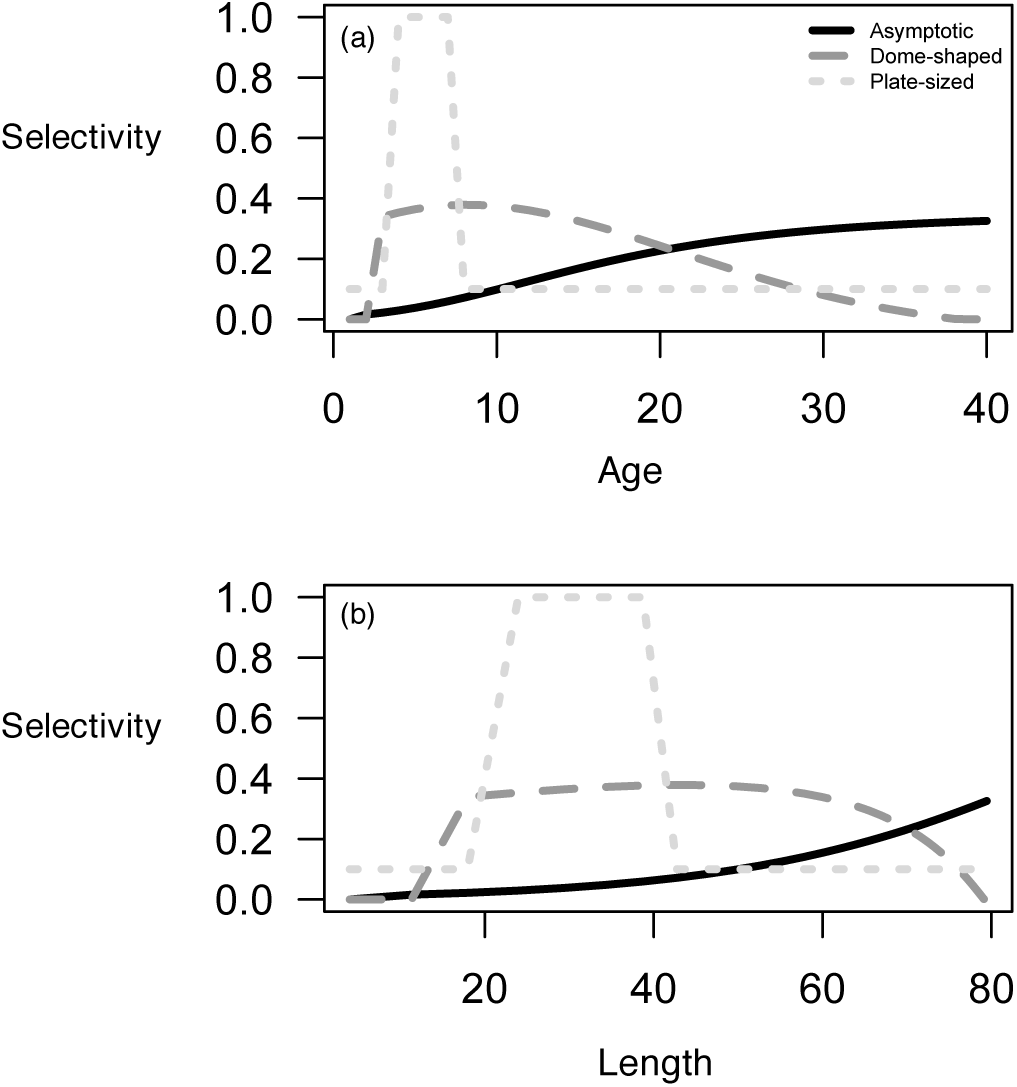
Asymptotic, dome-shaped, and plate-sized selectivity functions of (a) age and (b) length for the Brown-marbled grouper growth curve when *k* = 0.1. The area under each curve is equal. While the plate-sized curve reflects a strong preference for plated-sized fish, it includes some possibility of incidental capture or discard mortality of older or younger fish. Bottom panel: Selectivity as a function of size, not age.

We give the specific selectivity functions in Table 2. In each case, the fishing effort on each age or size class is a product of the selectivity and the maximum fishing effort possible, such that

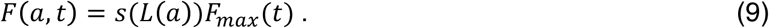

This approach to modelling selectivity differs from the approach taken elsewhere (Gwinn et al. 2015), where differences in selectivity (or vulnerability) are assumed to reflect different management tactics such as slot limits that have been put in place to control fishing effort. In other words, in our models a difference in selectivity equates to a difference in average fishing mortality across all ages. However, a change in growth patterns will also change the selectivity for each age, so that a change in growth rate may interact with selectivity to increase or decrease fishing mortality.

**Table 2.** Equations and parameters used to generate selectivity functions in Figure 2. Parameters are described in Table 1. Notice that asymptotic selectivity depends on length-at-age *L(a);* Dome-shaped selectivity follows a Weibull distribution. Parameters were chosen so that the total area under the curve was similar for each function when *k* = 0.1 and *L*_∞_ = 90 in order to ensure that fishing mortality was comparable over the lifetime in this baseline case.

| Selectivity | Function | Parameters |
| --- | --- | --- |
| Asymptotic probability | $s(L(a)) = \frac{1}{1 + e^{-j(L(a)-D)}}$ | $j = 0.05, D = 94$ |
| Dome-shaped probability | $s(a) = K + \left(\frac{x}{b}\right) \exp\left(-\left(\frac{x}{b}\right)^z\right)$<br>where $s(a)$ must be greater than 0 | $s(a) = 0$ for $a \leq 2$<br>$x_a = (a + 11)/12$<br>for $a > 2$<br>$K = -0.05, z = 2, b = 2$ |
| Plate-sized probability | $s(L) = \begin{cases} 0.1 & \text{if } L < L_{crit} \\ 1 & \text{if } L > L_{max} \end{cases}$ | $L_{crit} = 20$<br>$L_{max} = 40$ |

### Biological reference points and recovery metrics

We calculate two common biological reference points, as well as the maximum growth rate of the population during recovery after fishing stops. To do so, we denote the number of females in a given age class at the steady state by *N¯*(*a*); with fishing it is *N¯(a,F)*. The egg production of these females with fishing mortality *F* relative to egg production in the unfished population (often called the Spawning Potential Ratio (SPR; Mangel et al. 2013) is

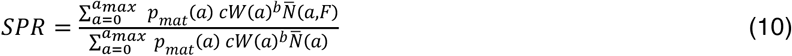

Note the similarity of the numerator and denominator of Eq. 10 to the egg production equation (Eq 5). For sex-changing stocks, the SPR is calculated in a similar fashion, but the analogous equation for egg production is Eq. 7. In both cases, the steady state biomass when fishing mortality is *F*, relative to the biomass of the unfished population is

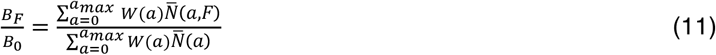

We also calculated the maximum growth rate of the recovering population, *λ_rec_*, after fishing (*F_max_* = 0.3) ends at *T_recovery_* (Fig. 3a). We determined *λ_rec_* by calculating the maximum difference in population biomass from one year to the next once fishing stopped, i.e.

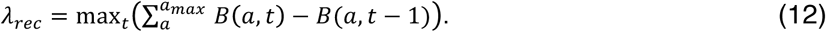

**Figure 3.**
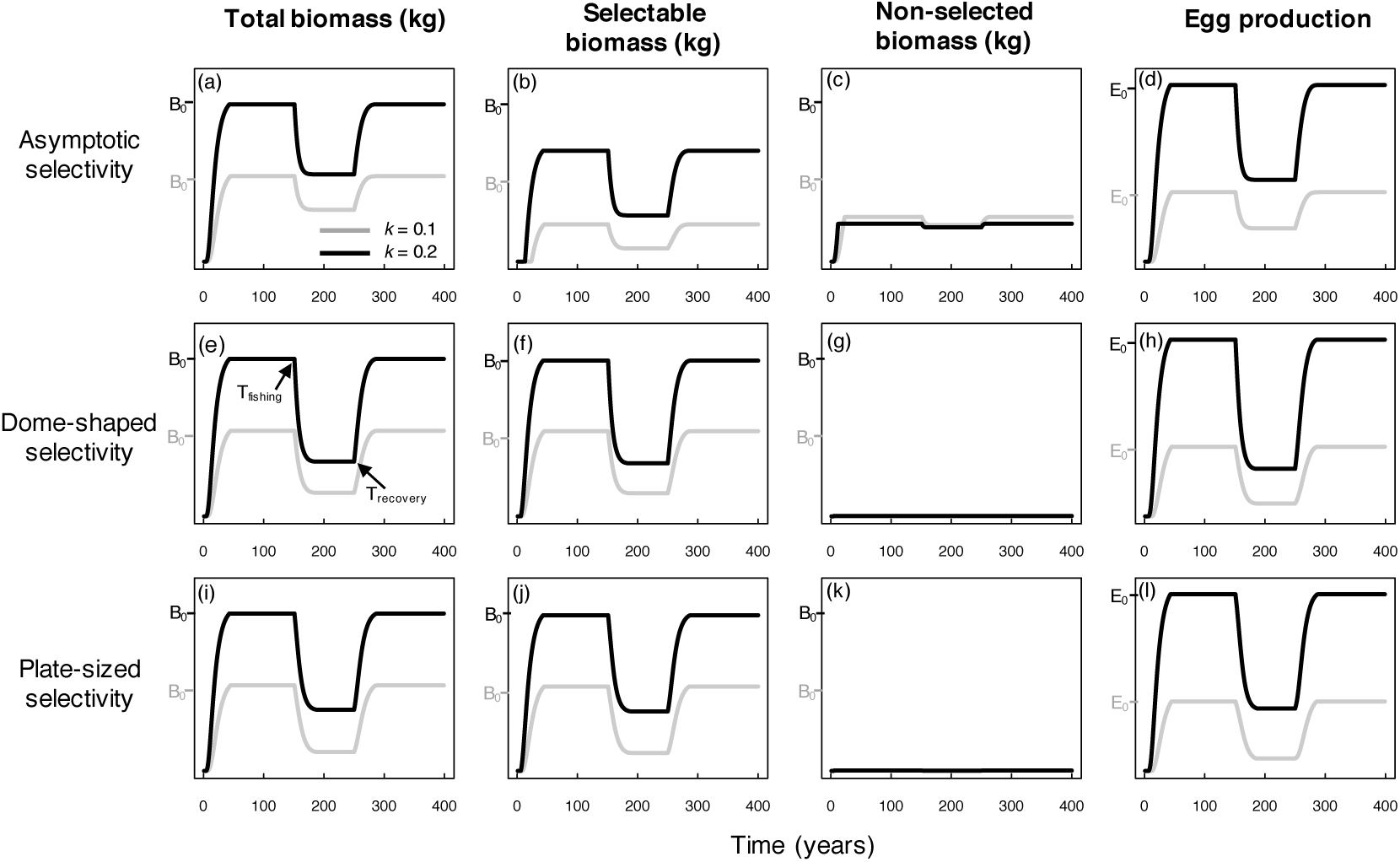
Comparison of biomass dynamics and egg production of the separate-sex version of the Brown-marbled grouper life history with three types of selectivity and two different growth coefficients, *k. F_max_* =0.3 and natural mortality *M*= 0.1.

## Results

The biomass dynamics of separate-sex and female-first stocks to fishing were similar for the Brown-marbled grouper-like life history. Plate-sized selectivity and dome-shaped selectivity, which select individuals near their age of maturation, decreased the steady-state biomass more than the asymptotic selectivity scenario that we considered (Fig. 3; only separate-sex biomass dynamics are shown). Stocks with slower somatic growth had lower selectable biomass (biomass caught with the combination of selectivity and fishing effort *F_max_*), and egg production in all three selectivity scenarios. With female-first sex change, egg production was always lower than in separate sex-stocks; the slow-growing female-first stocks had the lowest steady-state egg production of any population.

In Fig. 4, we show how fishing affects age-structure and sex ratio for both sexual systems (separate-sex, or gonochoristic, and female-first, or protogynous). In the separate-sex case, plate-sized selectivity eroded the population age structure most dramatically (Fig. 4). In the female-first population, due to plasticity in sex change, females began their transition to males earlier for all selectivities, and the average size of spawning individuals decreased (Fig 4). However, whether fishing affected female or male age structure most depended on the selectivity, the natural mortality rate (Table 3), and the shape of the sex change function.

**Figure 4.**
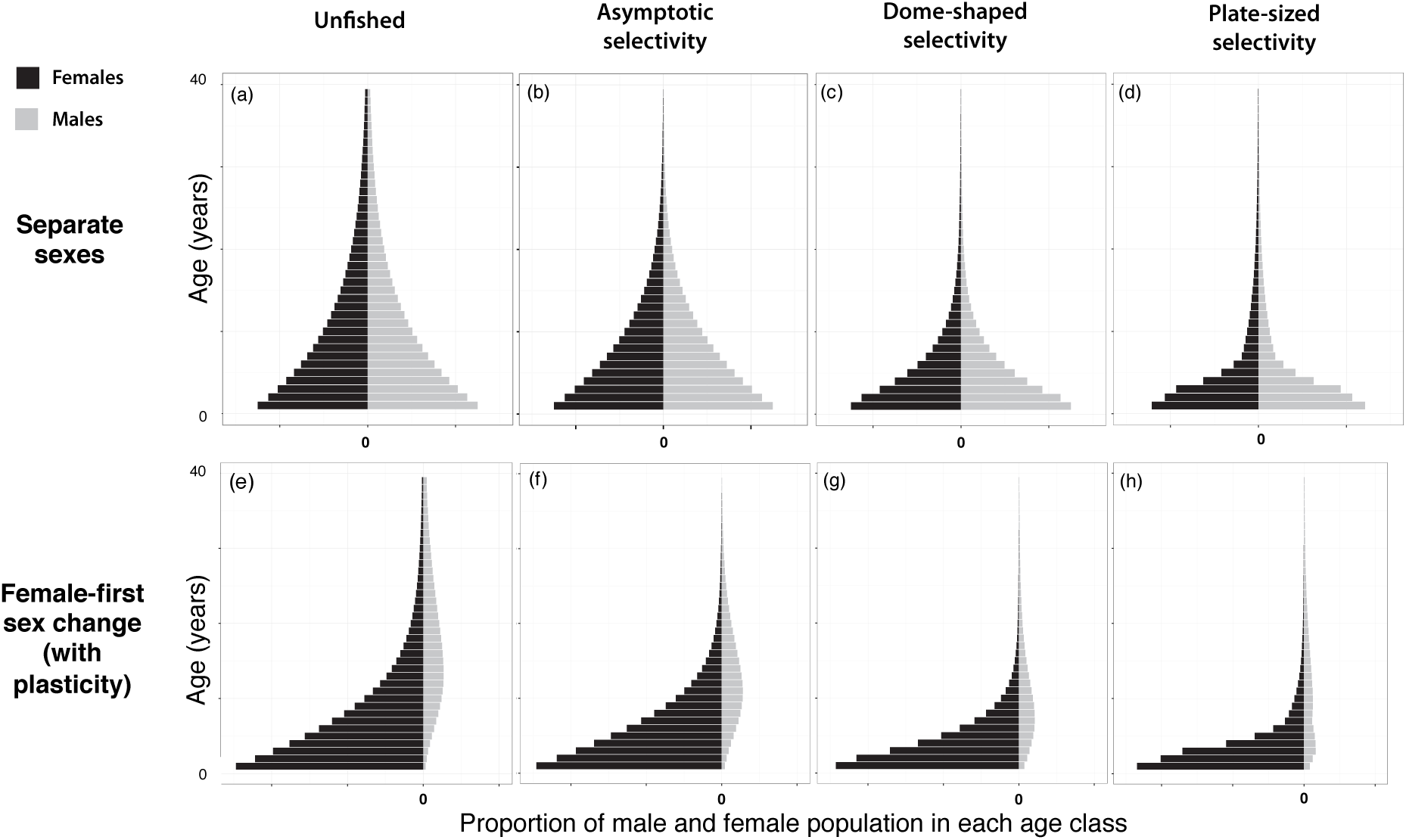
Change in age structure (the stable age distribution) for separate-sex and female-first stocks with different selectivity functions. Note that here we have assumed constant recruitment. Life history functions are from Brown-marbled grouper (left panels of Fig. 1), with parameters *L*∞ = 80 cm, *k* = 0.1, *M* = 0.1, and *F_max_* = 0.3, and *L_mat_* = 45 cm. Selectivity functions are in Fig. 2. The separate-sex stock (a-d) has symmetrical age structure for males and females because we assume a 50:50 sex ratio from birth. Sex-changing stocks (e-h) adjust the size at sex change according to the mean length of individuals in the population; in the unfished population *L_c_* = 75 cm.

In Table 3, we give the two biological reference points and the rate of recovery for both sexual systems under the three forms of fishing selectivity (with the same fishing mortality *F_max_*), as well as sensitivity to variation in natural mortality and somatic growth rate. The fishery with asymptotic selectivity had the greatest relative biomass at both levels of natural mortality and for both sexual systems (Table 3). Fishing had a greater relative impact on biomass (i.e., B_f_/B_0_ was lower) when natural mortality was lower for all selectivity functions. The biomass reference point (B_f_/B_0_) and the rate of recovery (*λ_rec_*) did not vary with sexual system in any of the three selectivity scenarios. However, the Spawning Potential Ratio (SPR) was always lower with female-first sex change than with separate sexes. Where biomass was depleted the most (i.e., with plate-sized selectivity and slow growth), populations had the highest rate of recovery, as this metric is greatest at low density. Finally, whether the SPR was lower with dome-shaped or plate-sized selectivity depended on both the background mortality rate and the slope of the sex change function. For example, in Table 3 the SPR with dome-shaped selectivity and *M* = 0.1 is lower than the SPR with plate-sized selectivity; this reverses with *M* = 0.3 because of the relationship between the average size of individuals in the population and their probability of transitioning from a female to a male.

**Table 3.**
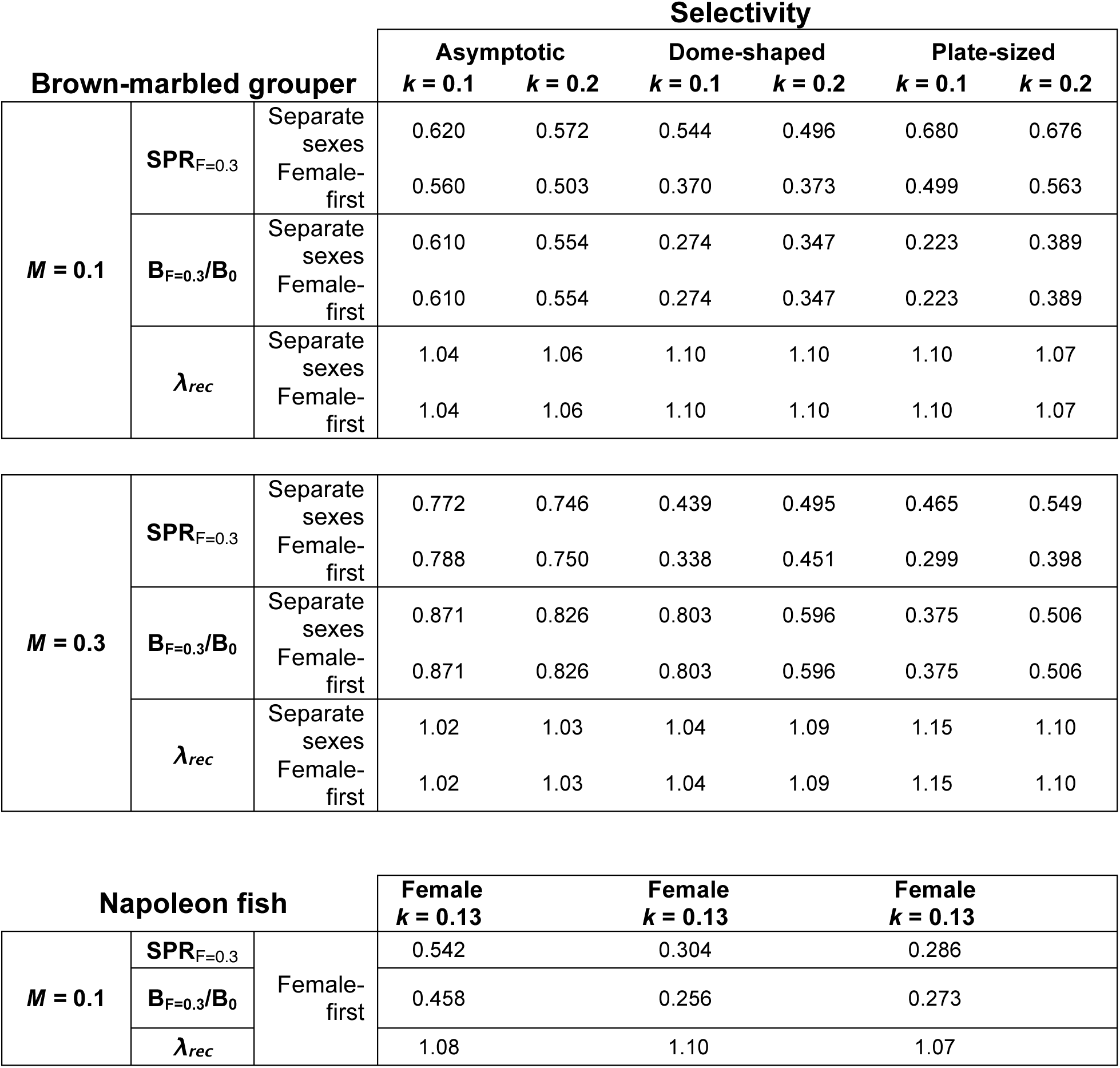
Sensitivity of population metrics (SPR_F_=_0.3_, B_F_=_0.3_/B_0_, and *λ_rec_* to variation in natural mortality rate *M* and growth rate *k* for each type of fishery selectivity and each example species. These reference points do not represent absolute differences in unfished biomass. For all cases, *F_max_* = 0.3. In the two upper panels, we calculate each metric for both mating systems and a life history based on Brown-marbled grouper. Notice that mating system affects SPR, but not relative biomass nor population recovery rate for Brown-marbled grouper. In the bottom panel, we calculate each metric for the life history based on Napoleon fish with no plasticity in the sex change function (also see Fig. S2).

| Brown-marbled grouper |  |  | Selectivity |  |  |  |  |  |
| --- | --- | --- | --- | --- | --- | --- | --- | --- |
|  |  |  | Asymptotic |  | Dome-shaped |  | Plate-sized |  |
| | | | $k = 0.1$ | $k = 0.2$ | $k = 0.1$ | $k = 0.2$ | $k = 0.1$ | $k = 0.2$ |
| $M = 0.1$ | $SPR_{F=0.3}$ | Separate sexes | 0.620 | 0.572 | 0.544 | 0.496 | 0.680 | 0.676 |
|  |  | Female-first | 0.560 | 0.503 | 0.370 | 0.373 | 0.499 | 0.563 |
| | $B_{F=0.3}/B_0$ | Separate sexes | 0.610 | 0.554 | 0.274 | 0.347 | 0.223 | 0.389 |
|  |  | Female-first | 0.610 | 0.554 | 0.274 | 0.347 | 0.223 | 0.389 |
| | $\lambda_{rec}$ | Separate sexes | 1.04 | 1.06 | 1.10 | 1.10 | 1.10 | 1.07 |
|  |  | Female-first | 1.04 | 1.06 | 1.10 | 1.10 | 1.10 | 1.07 |
| $M = 0.3$ | $SPR_{F=0.3}$ | Separate sexes | 0.772 | 0.746 | 0.439 | 0.495 | 0.465 | 0.549 |
|  |  | Female-first | 0.788 | 0.750 | 0.338 | 0.451 | 0.299 | 0.398 |
| | $B_{F=0.3}/B_0$ | Separate sexes | 0.871 | 0.826 | 0.803 | 0.596 | 0.375 | 0.506 |
|  |  | Female-first | 0.871 | 0.826 | 0.803 | 0.596 | 0.375 | 0.506 |
| | $\lambda_{rec}$ | Separate sexes | 1.02 | 1.03 | 1.04 | 1.09 | 1.15 | 1.10 |
|  |  | Female-first | 1.02 | 1.03 | 1.04 | 1.09 | 1.15 | 1.10 |

| Napoleon fish | | | Female<br>$k = 0.13$ | Female<br>$k = 0.13$ | Female<br>$k = 0.13$ |
| --- | --- | --- | --- | --- | --- |
| $M = 0.1$ | $SPR_{F=0.3}$ | Female-first | 0.542 | 0.304 | 0.286 |
| | $B_{F=0.3}/B_0$ | | 0.458 | 0.256 | 0.273 |
| | $\lambda_{rec}$ | | 1.08 | 1.10 | 1.07 |

In Figures 5 and 6 we show the effects of varying levels of fishing mortality on the SPR and steady state biomass (B_F_/B_0_) for each of the three forms of selectivity and for both sexual systems. Lower values of both reference points with plate-sized selectivity and dome-shaped selectivity hold across a range of fishing mortalities for both separate-sex scenarios (Fig. 5) and female-first scenarios (Fig. 6). Our asymptotic selectivity scenario had less of an effect on the reference points because the fishery selectivity was lower for adult age classes (even with large values of *F_max_*). When selectivity depends on size-at-age (as in our asymptotic selectivity function), growth rate had a surprising effect on the dynamics of biomass. Stocks with lower somatic growth rates had the greatest B_f_/B_0_ (Figs. 5d and 6d, Table 3), and these stocks declined more slowly than fast-growers as *F_max_* increased. This is because fast-growing fish entered the fishery earlier in life with this selectivity curve and therefore had higher catchable biomass. In other words, in fast-growers a larger proportion of the biomass was susceptible to capture (Fig. 3b).

**Figure 5.**
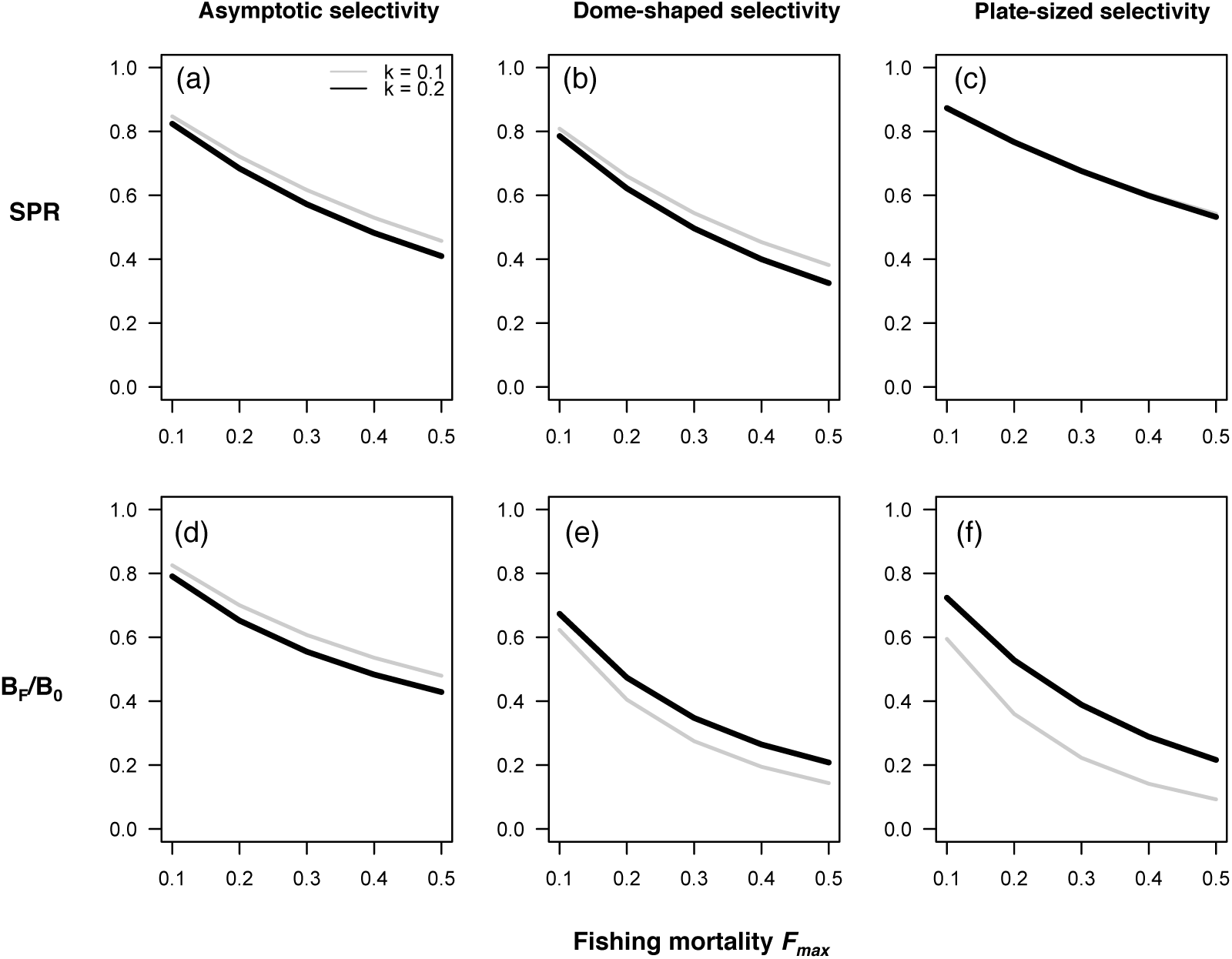
Sensitivity of biological reference points to fishing mortality for two growth rates in the separate-sex version of a Brown-marbled grouper-like life history. Each stock is fished with the three different selectivities in Fig. 2. (a-c): Spawning Potential Ratio (SPR), (d-f) Relative biomass (B_f_/B_0_). Natural mortality *M* = 0.1

The separate-sex stock with asymptotic selectivity (Fig. 5a) had a higher SPR with slow growth (Fig. 5a, Table 3) for a given mortality rate. In this case, the production lost from the delayed maturation of slow-growing females was compensated by females spending less time in susceptible size classes; this pattern was robust in our sensitivity analyses. We also found the separate-sex stocks had a higher SPR with dome-shaped and plate-sized selectivity if natural mortality was lower (Table 3). The unfished population biomass was lower with slow growth, suggesting that the relative effect of fishing mortality on productivity will be sensitive to both natural mortality and growth rate. By contrast, in female-first populations, slow-growing stocks always had a lower SPR (Fig. 6a-c) *except* when we investigated the scenario with age-dependent mortality and asymptotic selectivity (Fig. S3); here the SPR was higher for stocks with slow growth (comparable to separate-sex stocks). This was not true for the dome-shaped or plate-sized selectivity scenarios with age-dependent mortality.

**Figure 6.**
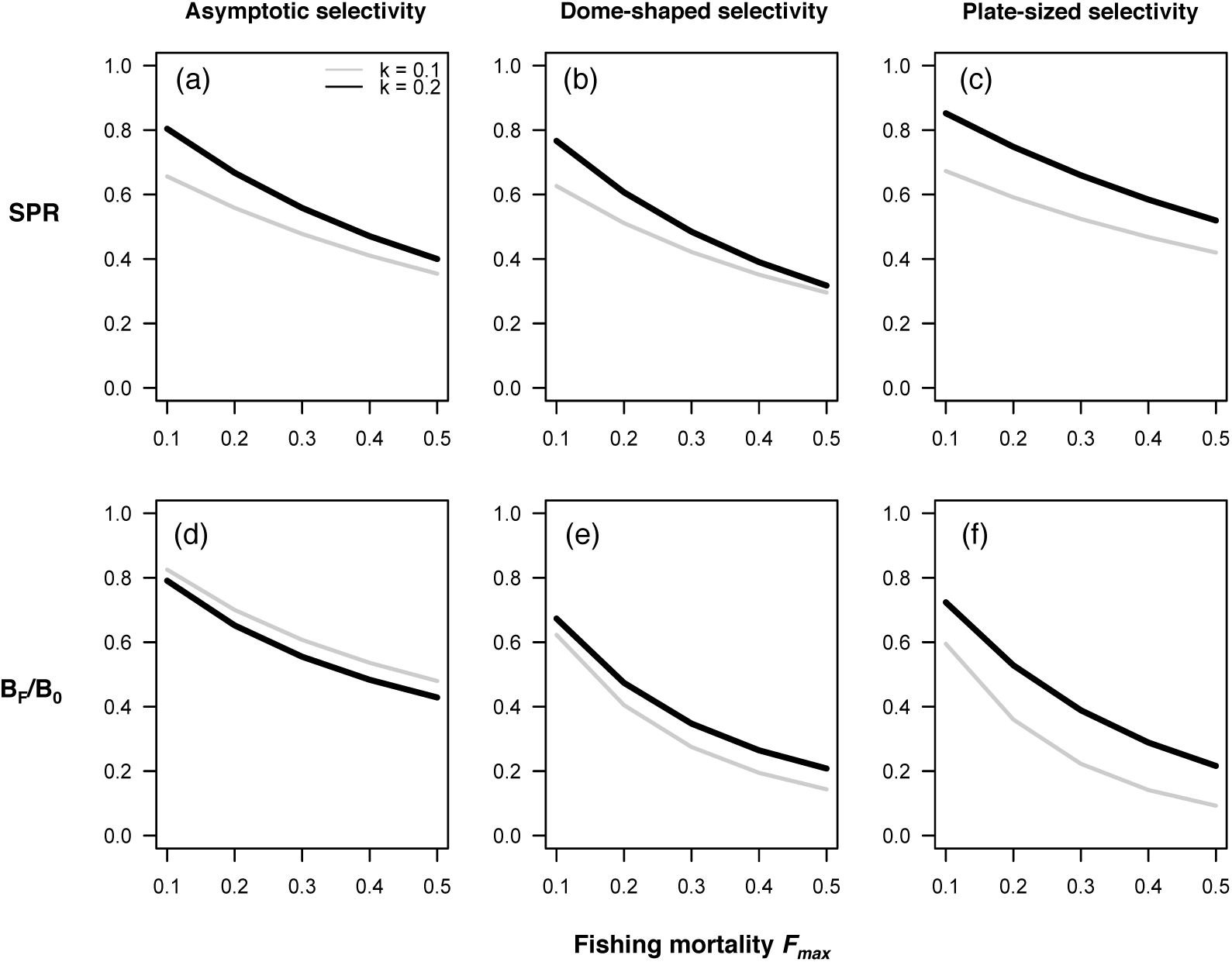
Sensitivity of two biological reference points to fishing mortality for two growth rates in the female-first Brown-marbled grouper-like life history. Each stock is fished with the three different selectivities in Fig. 2. Each stock is fished with the three different selectivities in Fig. 2. (a-c): Spawning Potential Ratio (SPR), (d-f) Relative biomass (B_F_/B_0_). Natural mortality *M* = 0.1.

In Figs. S1-S3 we explored the assumption of decreasing mortality with age in more detail. In this scenario, the unfished biomass and egg production increased for both separate-sex and female-first populations (Fig. S1), but the relative effects of fishing on the SPR and biomass (the biomass reference point) were identical to the case with constant natural mortality (Fig. 5). For slow-growing sex changers, the plate-sized fishery decreased absolute egg production the most (Fig. S1h). In this scenario, declining mortality with age primarily benefited male survival, especially as fishing reduced the average size of individuals and females became rare (Fig. S2). Hence the SPR was lowest for sex-changers with age-dependent mortality, and there was an interaction between selectivity and growth rate (Fig. S3). These results imply that the SPR is strongly affected by somatic growth rate, in addition to the details of natural mortality, sex change rules, and fishery selectivity and effort.

Until now, we have considered sensitivity analyses of the life history traits of the Brown-marbled grouper. We next compare these patterns with the results of the model developed to match the life history of Napoleon fish. This protogynous species differs from Brown-marbled grouper in that males reach much greater maximum sizes, though males and females have similar lifespans (Choat et al. 2006). Therefore, we modeled sex-specific growth rates (Fig. 1b). We also assumed the slope of the sex-change function for Napoleon fish was steeper than for Brown-marbled grouper (compare Fig. 1g and Fig. 1h), based on the apparent rarity of small males in the wild (Sadovy de Mitcheson et al. 2010). All life-history parameters are given in Table 1.

Whether or not we allowed for plasticity in the sex-change function for Napoleon fish had a minor effect on age structure after fishing (Fig. 7). Plasticity did not dramatically affect the biomass reference point, although the female biomass decreases more with plasticity, and the sex ratio became more biased without it (Fig. 7). Even without plasticity in sex change (which would decrease the female biomass further), the SPR of Napoleon fish was low for the dome-shaped and plate-sized selectivity scenarios (Table 3, Fig. S4). These selectivities targeted female biomass, thus reducing the compensatory capacity of Napoleon fish (Sadovy et al. 2003). Despite their lower SPR, the relative biomass (B_f_/B_0_) and recovery rates (λ_rec_) of Napoleon fish were largely similar to the results for protogynous Brown-marbled grouper (Table 3, Fig. S4). This is likely due to their larger body sizes overall.

**Figure 7.**
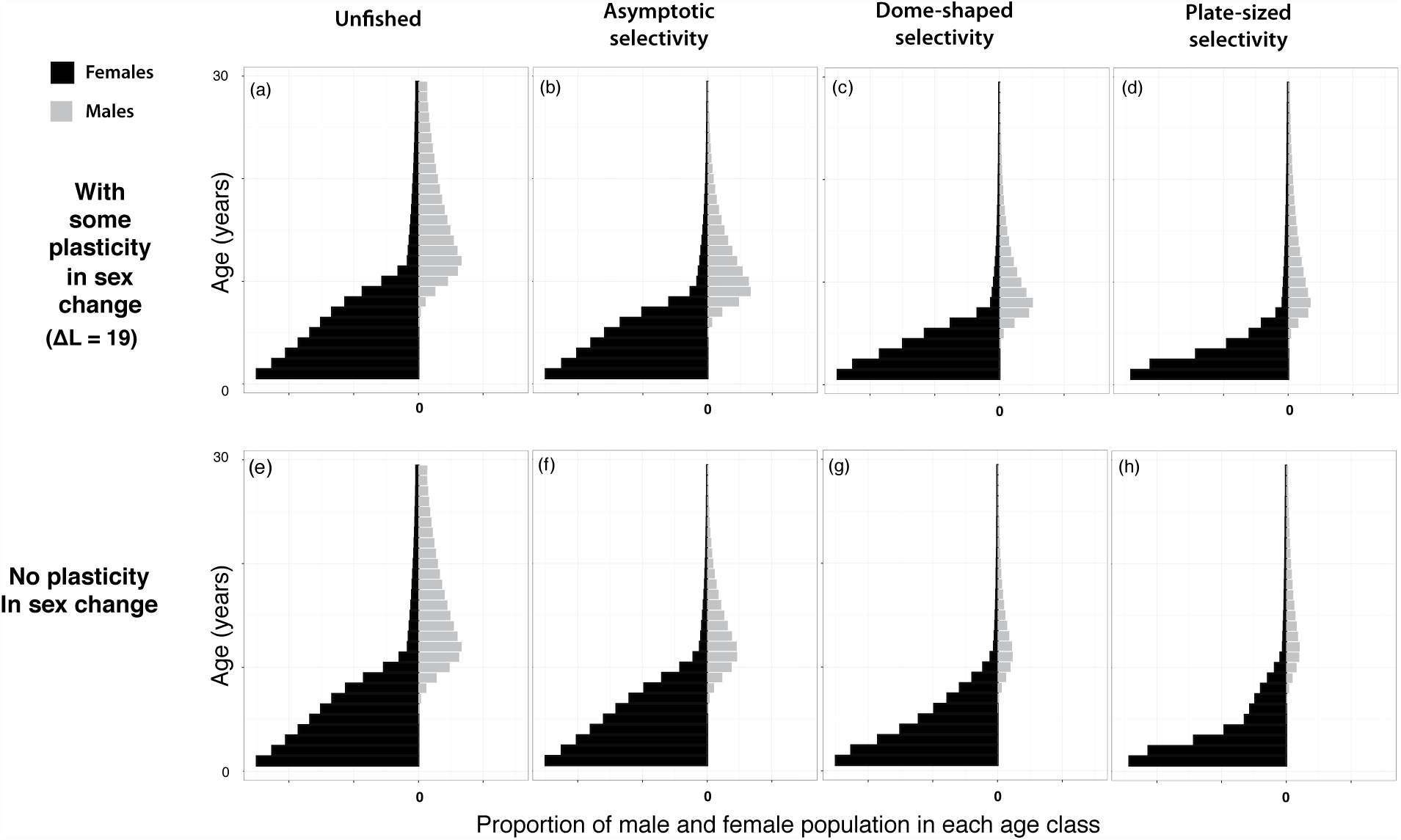
Stable age distribution (assuming constant recruitment) of Napoleon fish with (a-d) and without plasticity (e-h) in sex change. Females are black bars, and males are gray bars. Other parameters are in Table 1. Selectivity functions are in Fig. 2. Note that sex-changing stocks with some plasticity (a-d) adjust the size at sex change according to the mean length of individuals in the population.

## Discussion

Research on life-history correlates of species’ vulnerability to overfishing is often focused on fecundity, age at maturity, or body size (length or weight) (Reynolds et al. 2005, Patrick et al. 2009, Prince et al. 2015). Here, we have shown that somatic growth rate, maturation, and sex change interact differently with the three fishery selectivity scenarios considered. The aspects of selectivity that mattered most for sustainability are 1) the minimum age or size of capture relative to maturity and 2) whether selectivity depends on both age and size such that a change in growth rate changes the size-age relationship. Thus, for the scenarios we considered, plate-sized selectivity and dome-shaped selectivity had the greatest negative effect on population biomass (Fig. 3) and biological reference points (Figs. 5-6) because these fisheries targeted sub-adults that have already passed through the juvenile recruitment “bottleneck”, but had not yet begun to reproduce. Asymptotic selectivity had less effect because – as it targeted the greatest range of sizes – we assumed that the probability of being fished at any one size was lower overall in order to specifically compare selectivity effects when fishing mortality is similar across the lifetime.

Our choice of mass-specific fecundity parameters and the Beverton-Holt stock-recruitment relationship allowed populations to recover from low population sizes or extremely skewed sex ratios in our model. Even if the effects of fishing on age, size, and sex structure did not necessarily affect the biomass reference point (Table 3), the compensatory capacity of the stock always decreased. Specifically, the SPR decreased with *F_max_*, especially for hermaphroditic stocks where selectivity removed young fish (Table 3; Fig. 6 and Fig. S4), meaning that fishing decreased lifetime egg production the most in these scenarios.

Despite this difference in egg production, in our model there was no effect of sexual system on the biomass or the growth rate *λ_rec_* of recovering populations (Table 3). Were we to allow the possibility of Allee (depensatory) effects (such as a threshold density required for aggregation behavior or for fertilization of gametes), population recovery would be slowed (*cf.* Sadovy de Mitcheson 2016). Therefore, the insensitivity of biomass and recovery rate to protogyny applies only where aggregation behavior and fertilization of gametes is not impaired by changes in sex ratio or by severely reduced adult numbers. Furthermore, we did not address the possibility of episodic recruitment pulses, which are known to be important for the persistence of long-lived species (reviewed in Kindsvater et al. 2016). Given stochasticity in recruitment, we would expect that slow-growing protogynous species will be much more susceptible to overexploitation than populations or species with faster growth and separate sexes (i.e. gonochorism). Indeed, many populations of Napoleon fish have been severely depleted to the point where few recruits and young fish are observed (e.g. in parts of Indonesia, Sadovy de Mitcheson and Suharti unpublished), suggesting that stochastic recruitment limits the recovery of these populations. This process could be magnified by plasticity in the size-at-sex-change function because plasticity allows females to start transitioning to males at smaller sizes or younger ages (though this effect was minor in our analysis of Napoleon fish, given our assumptions). In species with plasticity in sex change, fishing decreases egg production in two possible ways: direct mortality increases, meaning that females are, on average, smaller, and also females transition to males earlier (see Vincent and Sadovy 1998). We suggest the effects of stochastic recruitment and its interaction with size- and sex-structure warrants further research.

Our results also highlight the effects of changes in growth rate for reference points and their interaction with selectivity. Fishing can affect growth rate directly, so that size-based selectivity can select for slow growth (Conover and Munch 2002). If fishing changes growth rates, we showed that the proportion of biomass vulnerable to fishing will also change if fishing depends on size-at-age, which might happen when time since birth affects probability of capture. In this case, in species or populations with faster growth, a greater proportion of the population will be vulnerable to fishing for the same selectivity. Evidence from Gag grouper – as well as meta-analyses across species – suggest that temperature and density both contribute to variation in the size-at-age relationship (Lindberg et al. 2006; Lorenzen 2008; Munch and Salinas 2011). The implications of this variation for fishery sustainability could be evaluated in future work.

Growth rate can also affect the relationship between size and sex change. Empirical evidence suggests that variation in growth due to latitude and temperature gradients, as well as habitat differences such as reef configurations, have strong effects on population biomass, density, and sex structure (Hamilton et al. 2007, Taylor 2014). In parrotfishes, e.g. Bullethead parrotfish (*Chlorurus spilurus*), variation due to environmental and social factors appears to swamp variation in vital rates due to fishing (Taylor 2014). In California Sheephead wrasse (*Semicossyphus pulcher)*, fishing decreases size and age at maturation and sex change (Hamilton et al. 2007). By contrast, in Hawaiian grouper (*Hyporthodus quernus*), there was little change in sizes of mature females over a 30-year period despite ongoing fishing, perhaps due to relatively strong fishery governance (DeMartini et al. 2011).

Prior models and reviews of the population dynamics of female-first hermaphrodites, including California Sheephead wrasse, Black sea bass (*Centropristis striata*) and Red hind (*E. guttatus*) have focused on sperm limitation due to observed effects of fishing on the population sex ratio with differential loss of males (Alonzo and Mangel 2004, Molloy et al. 2007, Brooks et al. 2008, Alonzo et al. 2008, Provost and Jensen 2015). For some hermaphroditic stocks, the number of males relative to females has decreased dramatically (reviewed in Shepherd et al. 2013). Similarly, in our model, female-first stocks declined to zero if selectivity drove the sex ratio below levels where all gametes were fertilized (not shown). This outcome was sensitive to the slope of the Beverton-Holt SRR near the origin and the shape of the sex change function (Fig. 1), which is consistent with previous models (Brooks et al. 2008). However, for some female-first sex-changing species, size-selective fishing is known to reduce the size at sex change (Hamilton et al. 2007, reviewed in Provost and Jensen 2015), although this evidence comes largely from studies of small reef fish (e.g., wrasses) (Taylor 2014). There are some protogynous species where the sex ratio has not changed with fishing, suggesting the sexchange rules are less plastic (Shepherd et al. 2013, Provost and Jensen 2015).

Our results add to this discussion by emphasizing that even if sperm limitation is not a factor (i.e., when sex ratios do not change), fishing can reduce egg production in female-first stocks. Limited egg production can reduce the ability of stocks to compensate for fishing or other disturbances (Kindsvater et al. 2016). Whether populations become egg-limited could depend on the mechanism and timing of sex change, but is most likely when fishing selectively removes more males. If some of the mature males are protected, females will be slower to transition to males (if sex change is plastic), and that could buffer the egg production of the population. This approach has been recommended elsewhere based on empirical observations (Pears et al. 2006, Williams et al. 2008).

### Sustainability of different capture methods for live reef food fish

We return now to our original motivation, understanding the effect of selectivity for “plate-sized” fish between 20 and 40 cm, as this practice is common for certain tropical groupers and wrasses targeted for the live reef food fish trade in Asia. As an alternative to the capture of plate-sized fish from wild populations, this size of fish can also be obtained by catching even smaller juveniles and growing them out in captivity to plate size. Both practices occur for several reef fishes, including Brown-marbled grouper and Napoleon fish. In both such cases, juveniles caught for “grow-out” are typically post-bottleneck and are raised in captivity for months or years. However, there are a few places where juveniles are taken at or very shortly after settlement and can be considered to be pre-bottleneck, meaning that fish at this stage have very high natural mortality levels. In one region of western Indonesia, fishers using hand nets take juvenile Napoleon fish as ‘settlers’ from inshore nursery areas. In this case the grow-out period after capture can take from 4-5 years (Y. Sadovy de Mitcheson pers. obs.).

While we did not explicitly model the capture of settlers, we can predict the effects of a grow-out fishery targeting newly settled juveniles (under 5 cm) on fishery sustainability. In our model, the density-dependent bottleneck before and during settlement is represented by the Beverton-Holt SRR (Eq. 8). When fishing occurs shortly following the age of settlement, the recruitment function can be modified to include fishing mortality (here noted as *F_settler_*). The Beverton-Holt SRR can be derived by following the cohort of eggs until they recruit to the population in which *m_1_* denotes the background mortality of juveniles, *m_2_* the strength of density-dependent compensation, and *τ* the time spent in the pre-recruit stage (Mangel 2006). The number of recruits *R* is then

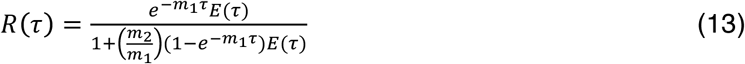

When we add fishing mortality in the early life history stage, the equation becomes

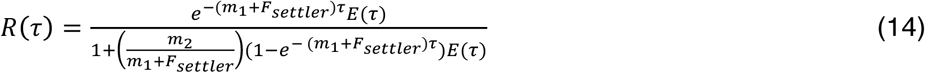

With this parameterization of the SRR, we see that both the numerator and the left-hand term in the denominator will decrease as fishing mortality *F_settler_* increases. This implies that targeting fish at or very near the age of settlement – when they naturally experience high natural mortality levels (pre-bottleneck) – may not affect productivity of the population unless *F_settler_* is so large that compensatory productivity is no longer possible. Whether increasing *F_settler_* affects recruitment to the post-bottleneck population depends on the duration of the settlement period *τ*, as well as the relative strength of *m_1_* and *m_2_*, and is worthy of further theoretical and empirical investigation.

Even without detailed information on the parameters in Eq. 14, we can infer the grow-out method of obtaining plate-sized fish by capturing (pre-bottleneck) settlers and maintaining them in captivity is more likely to be sustainable, all else being equal, than capturing similar numbers of juveniles or sub-adults, because the latter have much lower natural mortality. For example, if fishing reduces competition for hiding spaces or food, the removal of settlers will be compensated for by the increased survival of others. This will not be true when fishing removes sub-adult fish *after* they have survived the density-dependent bottleneck, where the effect of fishing will resemble the effects of plate-sized selectivity in our model.

The grow-out fishery for settlers could potentially be a productive method for obtaining high-value fish for the live reef food fish market, but it would need to be managed effectively. Settlers must be grown out for a long period of time (requiring food, and labour for up to five years), while the mortality rate of fish in cages during this period or during the process of collection is unknown and could be cumulatively substantial. Moreover, collection of settlers by fishers from the limited macro-algal habitat during the short seasonal recruitment period is extremely intense and could potentially remove most or all settlers; habitat destruction from the intensive hand net collection method used is also a concern. Further data and modelling work are needed to evaluate the quantitative implications of this type of fishing and produce recommendations for sustainable fishing and grow-out practices.

Importantly, the sustainability of a fishery for settlers will also depend on whether there is effective management limiting fishing of larger fishes, including subadults, in the same population. Fisheries for both juveniles and adults can be sustained biologically as long as fishing effort is regulated so that not too many sub-adults or large adults are taken. An example of this type of management is the Rock lobster (*Panulirus cygnus*) fishery in Western Australia, where take of both juveniles and adults is sustainable (de Lestang et al. 2012). This approach may be preferred if the aim is to maximize profit (assuming plate-sized fish are more valuable per kilogram than smaller or larger fish, as in the live reef food fish trade). The sustainable catch limit will depend on the biology of the species. For example, in Fig. 6, we showed that the sex-changing species subject to plate-sized-selectivity was still able to maintain a steady state, assuming there were no depensatory effects.

In addition to management that specifically addresses fisheries for sub-mature and pre-bottleneck fish, other management measures such as size limits (imposed by gear, trade or export restrictions) or temporal measures can be effective for data-poor fisheries (Dowling et al. 2008). In the case of aggregating species, temporal and spatial measures to protect spawning aggregations can restore stock age, size and sex structure (e.g. Molloy et al. 2009, Easter and White 2016). Temporal or spatial protection of spawning aggregations is likely to have a strong positive effect on population sustainability, as fish are especially vulnerable when aggregating (Erisman et al. 2015, Sadovy de Mitcheson 2016). However, theoretical work has cautioned that solely protecting a single population may have limited outcomes for other populations if fishing intensifies outside the protected area (Mangel 2000), if negative density-dependence limits growth and productivity of the protected population (Ellis and Powers 2012), or if the area is too small to protect enough adults adequately when not aggregating or does not encompass migration routes. This caveat is relevant to species where sex change is under social control, such that growth and sex change are related to the local density and size structure of mature fish. Our results suggest that size limits or spatial protection will be most effective if these measures are also designed to protect sub-mature individuals.

## Conclusions: guidelines for conservation and management

In the case of tropical fisheries there is an urgent need for general management guidelines that can be made with limited data and modelling work (Gruss et al. 2014, Prince et al. 2015).

**Table 4.** Sustainability of management guidelines for each type of fishery selectivity, given its interaction with sexual pattern; protogynous hermaphroditism versus gonochorism.

| **Fishery type** | **Separate sex (gonochoristic)** | **Female-first sex change (protogyny)** |
| --- | --- | --- |
| **Asymptotic selectivity** | Can be sustainable if the minimum length limit protects an adequate portion of the spawning stock and reproduction (i.e., aggregation) is possible | Can be sustainable especially if some of the largest males are protected to preserve social dominance hierarchies |
| **Dome-shaped selectivity** | May be sustainable if sub-mature fish are protected (as well as the largest fish), akin to a slot harvest management limit |  |
| **Selectivity for sub-mature fish (grow out or "plate-sized" fishery)** | A sustainable fishery is possible if managed to ensure low levels of catch biomass. Gear restrictions that protect fish up to sexual maturation could improve sustainability | A sustainable fishery is possible at very low catch biomass; otherwise fishing will erode productivity of the population, especially with skewed sex ratios |
| **Capture and grow out at settlement (pre-bottleneck selectivity)** | A sustainable fishery is possible if adult catch is also managed, cage mortality is low, and settlement habitat is protected |  |

Our results lead to several rules of thumb relevant for management in fisheries where specific size ranges of fish are targeted to meet market demand (Table 4). When selectivity targets fish prior to maturity, but after the initial high mortality experienced by juveniles at settlement (i.e., post bottleneck), it causes the steepest declines in biomass and productivity. Gear restrictions that protect fish up until the age of maturity could improve sustainability of the fishery.

1. Fishing erodes the compensatory capacity of slow-growing, female-first populations more quickly than fast-growing populations with separate sexes. Therefore, large protogynous groupers and wrasses with slow somatic growth are especially sensitive to overfishing and other environmental disturbances. This change in compensatory capacity (SPR) can happen before a decline in biomass is observed (Figs. 5 and 6).
2. Sustainable fisheries of plate-sized (sub-mature) fish are possible with strong enforcement of management limiting catch of juveniles and adults. If well managed, these fisheries could be more lucrative than fisheries for larger fish (despite being less productive biologically) in markets where plate-sized fish are more valuable per kilogram.
3. Fishing settlers (fish under approximately 5 cm in total length and pre-bottleneck) is potentially a productive source of fish and can be sustainable if managed. This also depends on fishery for larger individuals in the same population also being effectively managed (by suitable spatial, temporal, effort and/or gear controls) to maintain reproductive capacity. Settlement and nursery areas must be available, and mortality in captivity (during the grow-out phase) must be low; further data and simulation studies are needed to evaluate these processes.

## Acknowledgements

HKK was supported by NSF awards DBI 13-05929 and DEB 15-56779. JDR was supported by an NSERC Discovery Grant, and MM was funded by NSF awards OCE 11-30483, DEB 14-51931, and DEB 15-55729. We thank two anonymous reviewers for their comments, which improved an earlier draft of this manuscript.

